# Learning complex temporal dependencies via local synaptic plasticity

**DOI:** 10.64898/2026.07.09.737423

**Authors:** Julian Ng-Kee-Kwong, Mufeng Tang, Thomas Akam, Rafal Bogacz

## Abstract

The ability to extract and exploit temporal structure across diverse tasks is central to human cognition. Neuroscientists have typically relied on recurrent neural networks (RNNs) trained with backpropagation through time (BPTT) when modelling neural and behavioural processes such as decision-making and motor control. However, this algorithm has limited biological plausibility, hence the computational principles underlying efficient learning of temporal dependencies remain unresolved. Here, we investigate temporal predictive coding (tPC), a recently proposed framework that extends predictive coding to the temporal domain while preserving local Hebbian update rules. We analyse and extend tPC to establish its relationship with several influential computational models of learning in RNNs, including BPTT, reservoir computing, and eligibility propagation (e-prop). We first demonstrate structurally equivalent weight update rules between tPC and tBPTT1, a variant of BPTT in which gradients are propagated only one time step into the past. We then show that tPC can leverage reservoir dynamics to encode short-range temporal context, and simultaneously sculpt neural trajectories in state space to support downstream readout. We further demonstrate that hierarchical recurrent dynamics can facilitate learning of more complex temporal dependencies, while additionally conferring robustness to strong distractors. Finally, we show that tPC networks can be augmented with biologically inspired eligibility traces to solve temporally extended context-dependent tasks. Together, these results reveal that relatively simple recurrent networks governed by local plasticity can support temporal learning in more complex settings than previously appreciated.

**Author summary:** To navigate the world, our brains must constantly track how events unfold over time, whether we are predicting the next word in a sentence or timing a tennis swing. Neuroscientists often use artificial neural networks to study how the brain learns these sequences. However, these models are often not biologically realistic, since they adjust neuronal connections using information that individual neurons would not have access to in the brain. Here, we examine an alternative framework called temporal predictive coding, which is designed to better reflect how brain networks may be organised. We show that these biologically inspired networks are surprisingly powerful: they can remember recent context and, when arranged in layers, can learn increasingly complex patterns. We also show that they work better when equipped with a memory-like mechanism that helps neurons link recent events across time. Overall, our findings suggest that relatively simple, biologically inspired networks may capture how we process sequential events better than previously thought.

## Introduction

Humans successfully exploit the myriad regularities present in the external world to adapt and generalise to novel environments. Such regularities are not only spatial but also temporal in nature, unfolding across multiple timescales and often requiring the integration of information over extended periods. From understanding speech and music to predicting the behaviour of others and navigating dynamic environments, the ability to extract and leverage temporal structure is fundamental to cognition. Recent work has begun to uncover how temporal information may be represented across distributed neural populations, including the discovery of sequential neural activity and temporal context representations in cortical and hippocampal circuits [1–3]. An expanding literature has also characterised neural responses to violations of temporal expectations, providing evidence for temporal prediction error signals in cortical circuits [4–7]. Yet, computational accounts that explain how the brain might learn such temporal dependencies remain relatively underdeveloped.

In computational neuroscience, a standard approach for modelling tasks with temporal dependencies, such as decision-making or motor control, involves recurrent neural networks (RNNs) trained using backpropagation through time (BPTT) [8, 9]. However, BPTT is generally considered biologically implausible and unlikely to be implemented by the brain [10]. In particular, BPTT requires the precise storage of past neural states and the backward propagation of error signals across time [11]. Nevertheless, owing to its effectiveness and versatility, BPTT is still the dominant training method for RNNs in computational neuroscience, despite the development of alternative approaches with varying degrees of biological plausibility.

More generally, learning and inference in the brain remain subjects of considerable interest, and a variety of explanatory frameworks have been proposed [12–15]. The theory of predictive coding is one such well-established framework, widely used by neuroscientists to explain perceptual processes [15–18]. Consistent with the Bayesian Brain hypothesis [19], predictive coding posits that the cortex constructs internal models of the external world and uses them to generate predictions about incoming sensory input. A key appeal of predictive coding is that it not only provides a conceptual account of perception in terms of prediction error minimisation, but also offers a biologically plausible neural implementation based on local learning rules. To date, however, most predictive coding formulations have focused on static settings in which the temporal structure of stimuli is not explicitly modelled [20, 21].

Consequently, predictive coding networks have recently been augmented with recurrent dynamics to enable temporal prediction [22]. This framework, termed temporal predictive coding (tPC), has been shown to approximate Kalman filtering in both linear and non-linear dynamical systems, while retaining simple update rules based on local Hebbian plasticity [22]. Moreoever, tPC has been found to outperform Hopfield networks on sequential memory tasks [23], and to develop visual and spatial representations resembling those observed in biological neurons [22, 24]. A natural next step is therefore to assess whether tPC could serve as an alternative to BPTT-trained RNNs for modelling tasks involving more complex temporal dependencies.

Here, we demonstrate how the tPC framework could be extended to solve several such tasks, without requiring explicit storage of past neural states or deviating from local plasticity rules. After establishing a structural equivalence between the weight updates of tPC and a truncated form of BPTT known as tBPTT1, we show that tPC can exploit its reservoir dynamics to encode temporal structure while simultaneously shaping neural trajectories to facilitate downstream readout. We further demonstrate how a hierarchical architecture could enable more efficient learning of non-trivial temporal dependencies and improve robustness against perturbations induced by strong distractors. Finally, we show how eligibility traces may be incorporated into the tPC framework to support learning over extended temporal horizons.

Altogether, although further progress may be required to match the performance of BPTT in large-scale settings, our results demonstrate that relatively simple recurrent networks with update rules governed by local Hebbian plasticity can nevertheless learn complex temporal structure. In doing so, this work also clarifies the relationship between tPC and several influential computational frameworks, including reservoir computing, BPTT, and eligibility propagation (e-prop). More broadly, these findings suggest that biologically plausible mechanisms may support temporal credit assignment across more complex settings than is typically recognised.

## Results

### Temporal Predictive Coding

We start by providing a brief overview of tPC in this section. A more comprehensive treatment, including full derivation, is provided in the original publication [22]. For an intuitive introduction to the broader free-energy principle, from which predictive coding may be derived, we refer the reader to a tutorial by one of the authors [25].

The tPC framework, which assumes a graphical structure similar to that of an RNN, models the world as a sequence of hidden states 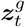 and a corresponding sequence of observations ***y***_*t*_, with optional external inputs ***x***_*t*_ (Fig 1a). Here, 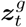 denotes the latent state posited by the generative model and is not directly observed. The model assumes that these latent states evolve over time and generate observations according to:

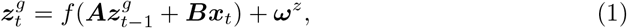

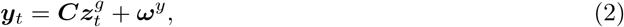

where ***A, B*** and ***C*** denote the recurrent, input and output weight matrices, respectively, and ***ω***^*z*^, ***ω***^*y*^ are sources of Gaussian noise. The function *f* may be an arbitrary non-linear activation function, here taken to be a tanh non-linearity. For simplicity, we do not consider an output non-linearity. Of note, although the original formulation applies the non-linearity prior to summation [22], we instead apply it after summation in order to facilitate direct comparison with standard RNN formulations in later sections. We use 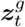 here to denote the underlying hidden state and will refer to its estimate simply as ***z***_*t*_. Since 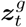 is unobserved, ***z***_*t*_ is the quantity used throughout the equations and computations that follow.

**Fig 1.**
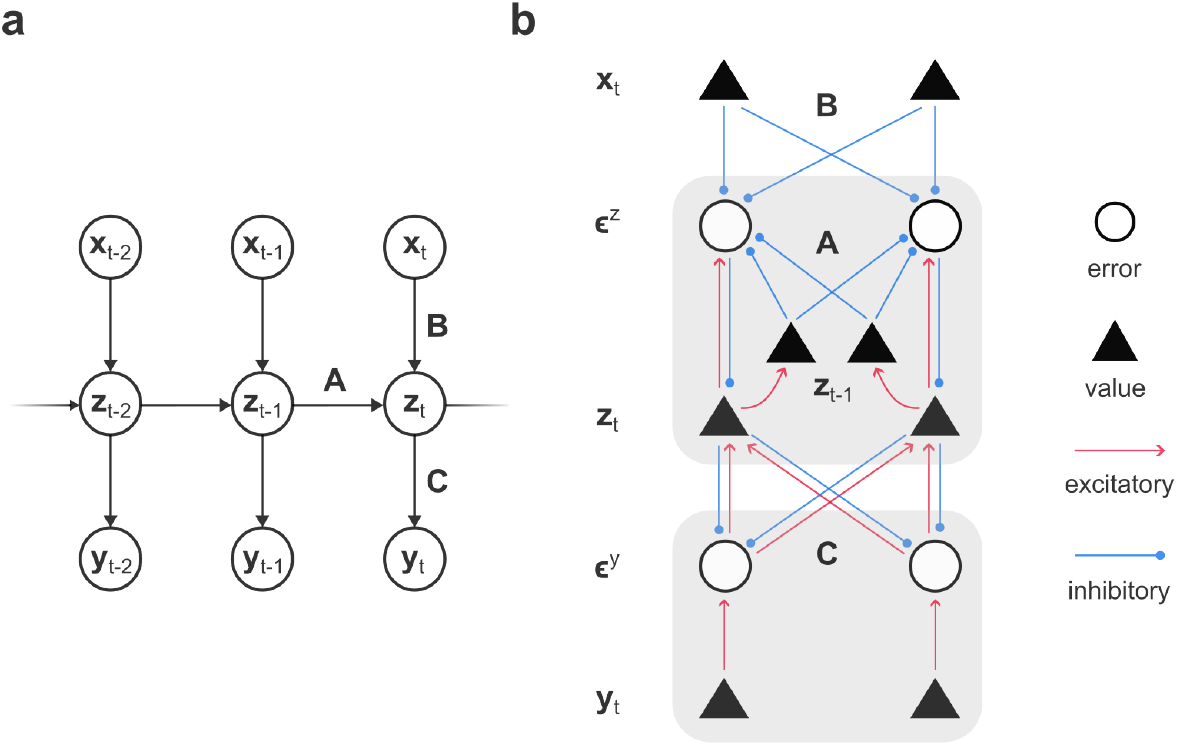
Temporal predictive coding model. **a:** Graphical model of tPC, in which the output ***y***_*t*_ depends on the hidden state ***z***_*t*_, which in turn depends on the input ***x***_*t*_ and the previous hidden state ***z***_*t*−1_. **b:** Possible circuit implementation of tPC using value and error units, which represent estimates of latent or observed variables and the discrepancy between neural activity and its top-down prediction, respectively.

### Objective

As with predictive coding frameworks, tPC may be formulated as a Bayesian inference problem where the aim is to infer the current hidden state 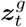 based on the previous estimate of the hidden state ***z***_*t*−1_, the current observation ***y***_*t*_ as well as the input ***x***_*t*_. Directly estimating the posterior distribution 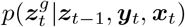 is however generally considered intractable. The free-energy principle therefore proposes a variational approach to Bayesian inference [15, 26]. The true posterior is approximated using a tractable distribution 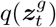, taken here to be a Gaussian under the Laplace approximation; we denote its mean (or equivalently, its mode) by ***z***_*t*_. Thus, 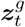 denotes the latent state of the generative process, whereas ***z***_*t*_ denotes the model’s inferred estimate of that state. Variational inference proceeds by minimising the variational free energy—a tractable objective whose minimisation also minimises the divergence between the approximate and true posteriors. Assuming ***ω***^*z*^, ***ω***^*y*^ are Gaussian with identity covariance, this quantity may be reduced, up to additive constants, to a sum of squared prediction errors:

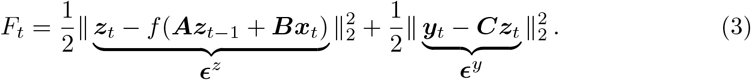

The first term corresponds to the difference between the inferred and predicted hidden state (‘temporal’ prediction error) while the second term corresponds to the difference between the true and predicted observation (‘sensory’ prediction error).

### Learning and inference

Minimisation of the free energy proceeds via alternation between gradient descent optimisation over the hidden state (i.e., inference) and over the weights (i.e., learning), analogous to expectation-maximisation (EM) algorithms. Differentiating the free energy with respect to the hidden state (and taking the negative) yields the update equation for inference:

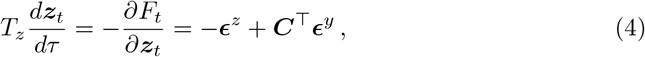

where *T*_*z*_ is the neuronal time constant. In practice, several steps of gradient descent are performed to allow inference to converge between observations of successive sensory inputs ***y***_*t*_—we have denoted this separation of timescales here via two distinct time indices, *t* and *τ*, for learning and inference, respectively. This may be reasonably justified on the basis that neural activity typically changes on a faster timescales than sensory inputs.

Taking the negative of the derivative of the free energy with respect to the weights leads to the update equations for learning:

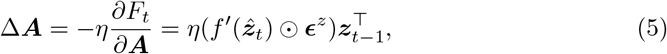

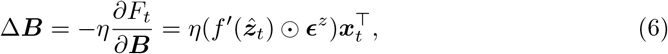

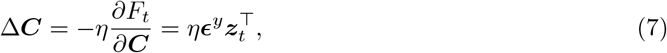

where

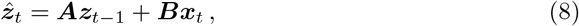

and *η* is the learning rate. *f*^*′*^(·) is the element-wise derivative of *f*, and ⊙ denotes element-wise multiplication.

Once trained, the model can generate predictions without performing inference by propagating activity forward using the learned weights:

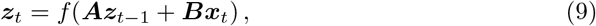

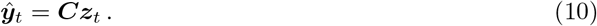

These equations correspond to the fixed point of the inference dynamics (Eq 4) when the output is unclamped, such that the sensory prediction error ***ϵ***^*y*^ is zero. In this setting, the hidden state is determined solely by the recurrent dynamics and current input, making prediction equivalent to the forward pass of a standard RNN with the same architecture.

### Neural implementation

tPC networks retain a similar architecture to standard predictive coding implementations (Fig 1b), wherein neurons are organised into two types: ‘value neurons’ which encode estimates of latent states or observations, and ‘error neurons’ which encode prediction errors. The input to value neurons in Fig 1b correspond to the right hand side of Eq 4, while the inputs to prediction error neurons correspond to their definitions in Eq 3. In tPC networks, prediction error neurons encode the difference between current activity and that predicted based on the input and the previously inferred state, stored in the additional neural population labelled ***z***_*t*−1_ in Fig 1b.

Under the proposed neural implementation, the weight updates defined in Eqs 5–7 are inherently local, depending only on quantities that are directly available at the relevant synapse. For example, changes in weights ***A*** depend on 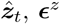 and 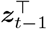 (Eq 5). Weights ***A*** describe connections between neurons encoding ***ϵ*** and 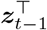, so these values are directly available at the synapses. Additionally, 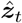 (defined in Eq 8) is a part of the input to neurons encoding ***ϵ***^*z*^ (defined in Eq 3), hence the neurons encoding ***ϵ***^*z*^ have access to 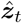. We refer the reader to the original publication for a detailed discussion on the possible neural implementation of tPC [22].

### Relationship between tPC and BPTT

Before empirically assessing the performance of tPC, we first clarify its relationship to BPTT, the standard algorithm for training RNNs. This section proceeds as follows: We begin with a brief exposition of BPTT [11] and review the tBPTT1 algorithm, in which gradients are not propagated beyond the previous time step. We show theoretically that tPC shares structurally equivalent weight update rules with tBPTT1, but differs importantly in how activity is updated at each time step due to the inference process. We additionally summarise under which conditions the tPC update rules end up converging to those of tBPTT1. We conclude by illustrating this distinction empirically on a position estimation task. For the theoretical analysis, we present the main results here and provide the relevant derivation in the Methods (subsections “Derivation of BPTT update rules” and “Derivations of tPC update rules and fixed point”).

### The BPTT algorithm

BPTT can be viewed as the temporal analogue of spatial backpropagation. It proceeds by unrolling the recurrent network across time and propagating gradients backward through the resulting computational graph. For an input sequence ***x***_*t*_ and a target sequence ***y***_*t*_ of length *T*, the mean squared error (MSE) loss evaluated at each time step is given by:

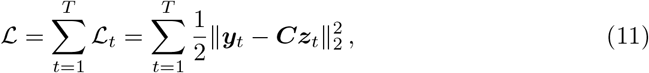

where we assume an RNN with forward dynamics given by

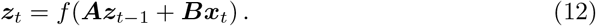

Of note, although the above equations closely align in mathematical structure with the deterministic components of Eqs 1 and 2, we wish to point out that the two sets of equations should be interpreted differently. For instance, Eq 12 defines the deterministic forward update of the RNN hidden state, while Eq 1 specifies the transition model of the tPC probabilistic generative model. Nonetheless, we use the same symbols to denote the corresponding variables of both models to emphasise their similarity, and clarify in the text which model the equations refer to.

The gradient descent update rule for the recurrent weight matrix ***A*** of an RNN can be written as (derived in Methods, subsection “Derivation of BPTT update rules”):

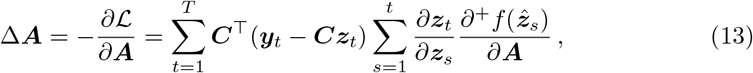

where 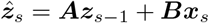. Since the recurrent weights are shared across time steps, each loss term in Eq 11 depends on ***A*** both through its immediate effect at time step *t* and through its earlier effects on hidden states that are propagated forward by the recurrent dynamics. The notation 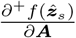 denotes the immediate partial derivative, in which the previous hidden state ***z***_*s*−1_ is treated as fixed with respect to ***A***. An analogous expression can be derived for the input weights ***B***.

BPTT is often truncated in practice to reduce computational cost and to mitigate vanishing or exploding gradients [27, 28]. In truncated BPTT, gradients are propagated backward only over a finite truncation horizon *K*, giving

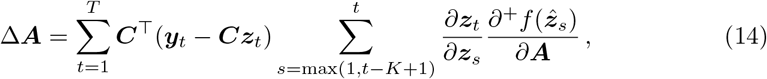

where the resulting update approximates the full BPTT update by retaining only temporal dependencies within the horizon *K*.

### tPC and tBPTT1 share structurally similar weight updates

The limiting case *K* = 1, which we refer to as tBPTT1, retains only the immediate dependence of the current hidden state on the recurrent weights:

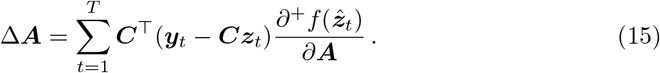

It is shown in the Methods (subsection “Derivations of tPC update rules and fixed point”) that, at convergence of the inference dynamics, the tPC update for the recurrent weights has an identical form:

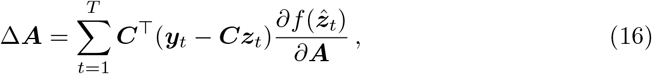

with the notational difference that, in the tPC expression, the final factor is not usually written explicitly as an immediate partial derivative. Thus, under these assumptions, the tPC update is equivalent in form to the tBPTT1 update; it captures the immediate effect of the recurrent weights on the current hidden state, but omits the additional temporal credit assignment terms that arise from propagating gradients through future hidden states.

### tPC inference results in different activity dynamics

However, because tPC includes an additional inference step updating ***z***_*t*_, an important difference arises at the level of activity updates. To illustrate this difference analytically, we consider the linear case. Although the tPC generative model assumes the same form of forward dynamics as a standard linear RNN, the resulting activity updates differ. In a standard RNN, the recurrent prediction directly determines the hidden state:

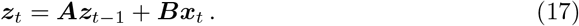

In tPC, by contrast, the corresponding prediction is refined through inference using the current observation ***y***_*t*_. Specifically, the fixed point of the tPC inference dynamics described by Eq 4 is given by (derived in Methods, subsection “Derivations of tPC update rules and fixed point”):

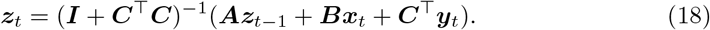

The inferred tPC hidden state therefore depends not only on the recurrent prediction, ***Az***_*t*−1_ + ***Bx***_*t*_, but also directly on the current observation through ***C***^⊤^***y***_*t*_, with (***I*** + ***C***^⊤^***C***)^−1^ acting as a gain term determining how these two sources of information are combined. Thus, although the tPC generative model and the RNN assume the same form of forward dynamics, tPC introduces an implicit flow of information from the observation to the hidden state at each time step, which is not present in a standard RNN trained with tBPTT1 (Fig 2a).

**Fig 2.**
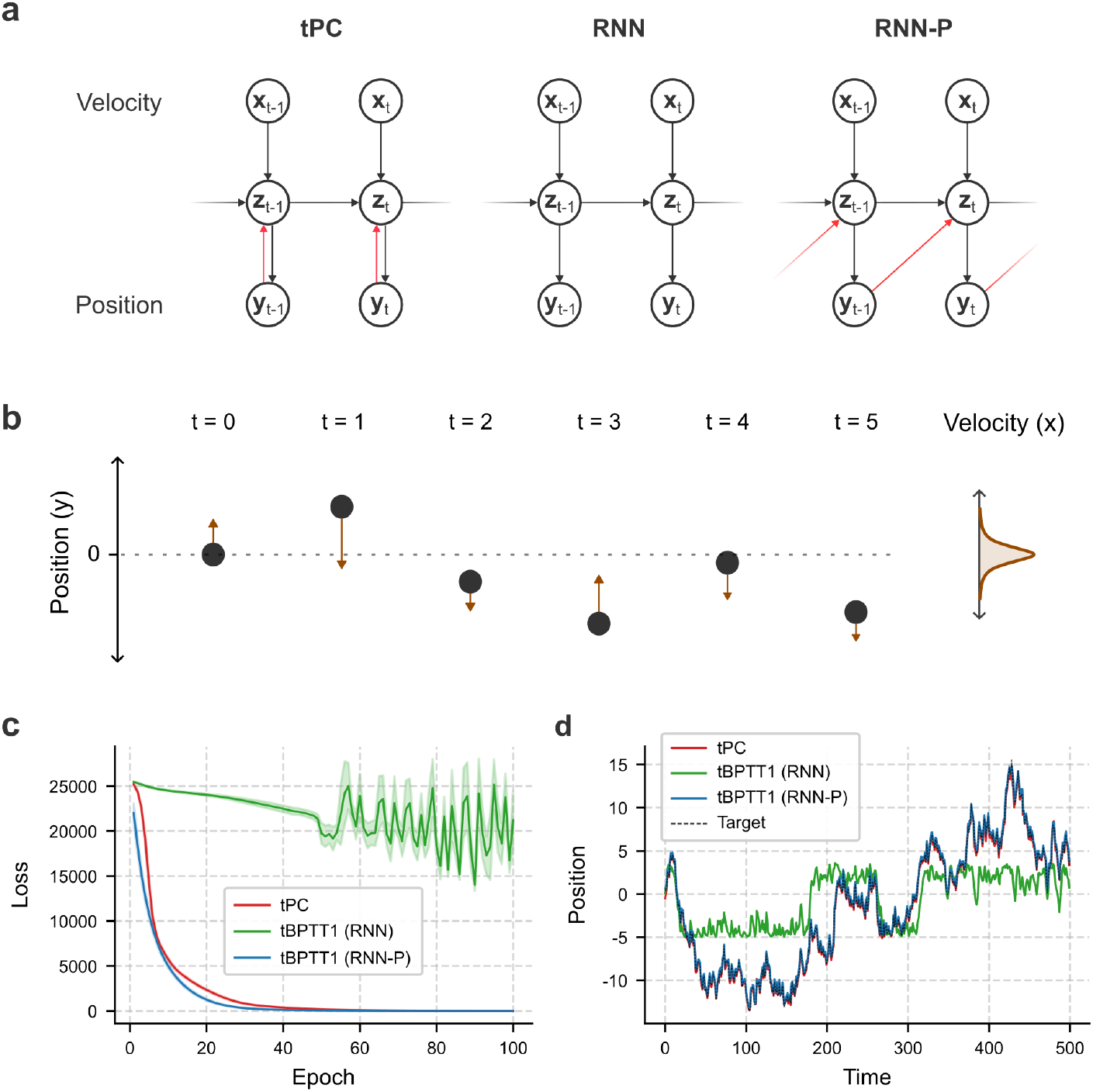
tPC inference induces implicit flow of information from observation to hidden state. **a:** Comparison of three models on the position estimation task: tPC, RNN, and RNN-P. Red arrows indicate flow of information from observation to hidden state. **b:** Schematic of the position estimation task, in which the model must predict the particle’s new position at each time step given a velocity input sampled from a standard Gaussian distribution (*µ* = 0, *σ* = 1). **c:** Training loss curves for the position estimation task, showing that a standard RNN trained using tBPTT1 fails to learn the task. **d:** Predicted particle trajectories for a representative sequence of velocity inputs, illustrating that both the tPC and RNN-P models can closely reproduce the target trajectory.

### tPC and tBPTT1 are equivalent under certain conditions

Before empirically examining the difference described above, we briefly consider the conditions under which tPC and tBPTT1 become equivalent, drawing on results relating predictive coding to backpropagation [29]. The key requirement for exact equivalence is that tPC inference does not alter the hidden state from that produced by the corresponding RNN forward pass. If the two models produce the same hidden activity at each time step—in the linear case, such that Eq 18 reduces to Eq 17—their corresponding weight updates also coincide (Eqs 15 and 16).

There are several cases in which the updates approach one another as the influence of the observation on tPC inference diminishes. First, when the output weights ***C*** are equal to zero, then Eq 18 reduces to Eq 17. Second, as the predicted observation ***ŷ***_*t*_ approaches the target ***y***_*t*_ during learning, the sensory prediction error ***ϵ***^*y*^ approaches zero, causing the inferred hidden state to approach the forward pass value. Third, a similar limit arises in the full formulation with explicit covariance terms (see [22]) when the output covariance becomes large relative to the covariance at the hidden layer, reducing the precision and therefore the influence of sensory prediction errors on inference. Under these conditions, predictive coding updates converge to those of backpropagation and, correspondingly, tPC updates converge to those of tBPTT1. In fact, analogous conditions can also be extended to hierarchical predictive coding architectures [29]. Although the exact equivalence in these special cases is of theoretical interest, we next show that tPC may differ from tBPTT1 in practice.

### Empirical results

Here, we demonstrate using a simple position estimation task (Fig 2b) that tPC and tBPTT1 may exhibit different behaviour due to the tPC inference dynamics resulting in information flow from observation to hidden state. In this task, a particle is initialised at position *y* = 0. At each time step, the model receives a velocity input sampled from a Gaussian distribution with mean 0 and standard deviation 1, and must predict the updated particle position. For a standard RNN, this corresponds to a path integration problem: the model must accumulate velocity inputs over time, starting from the initial position. By contrast, the task should be easier for tPC, because position information is implicitly available at each time step through the inference process. To isolate the contribution of this additional information, we compare tPC against an augmented RNN that receives the previous position as an explicit input alongside the velocity signal. This model, which we denote RNN-P (where P stands for previous), updates its hidden state according to:

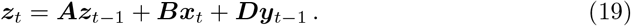

Consistent with this prediction, path integration is difficult over long trajectories for a tBPTT1-trained RNN (Fig 2c). However, when position information is available at each time step, as in tPC or RNN-P, the task can be solved much more easily. We briefly note that the performance of a variant of tPC, which receives the previous observation as explicit input (which we refer to as tPC-P), closely matched that of RNN-P (S1 Fig). These results suggest that tPC can be viewed as equivalent in form to tBPTT1 at the level of the weight update, while differing at the level of activity inference: tPC allows observations to influence the hidden state directly, thereby introducing an implicit observation-to-state information flow.

### Learning non-Markovian sequences via state-dependent computation

Building on the above theoretical results, which provide insight into tPC’s update rules, we next investigate the capacity of tPC to learn temporal dependencies. While we previously demonstrated that tPC can model simple dynamical systems using local Hebbian plasticity [22], here we extend this analysis to sequence prediction tasks involving longer and more complex temporal dependencies. Due to its close relationship to tBPTT1, it is unclear whether tPC can successfully model non-Markovian sequences, i.e., sequences where the observation at one time step depends not only on the previous observation. Here, we show this is possible by drawing from the framework of reservoir computing.

Reservoir computing, which was introduced in the early 2000s, allowed RNNs to process time-series data with minimal training cost [30, 31]. In reservoir computing, recurrent weights—and often the input weights as well—are kept fixed (Fig 3a). Instead, reservoir computing relies on non-linear dynamics to transform an incoming temporal signal into a high-dimensional representation. A simple, usually linear, readout layer may then be trained to map reservoir states to outputs. Popular implementations include the rate-based Echo State Networks (ESNs) and the spiking-based Liquid State Machines (LSMs), both of which exploit the transient dynamics and fading memory of recurrent systems to encode temporal structure in sequential data. In other words, since past inputs persist in the evolving internal state, the trained readout can infer temporal context without explicitly storing past observations. We reasoned that this ‘reservoir effect’ must also be inherent to tPC, given its similarly recurrent architecture.

**Fig 3.**
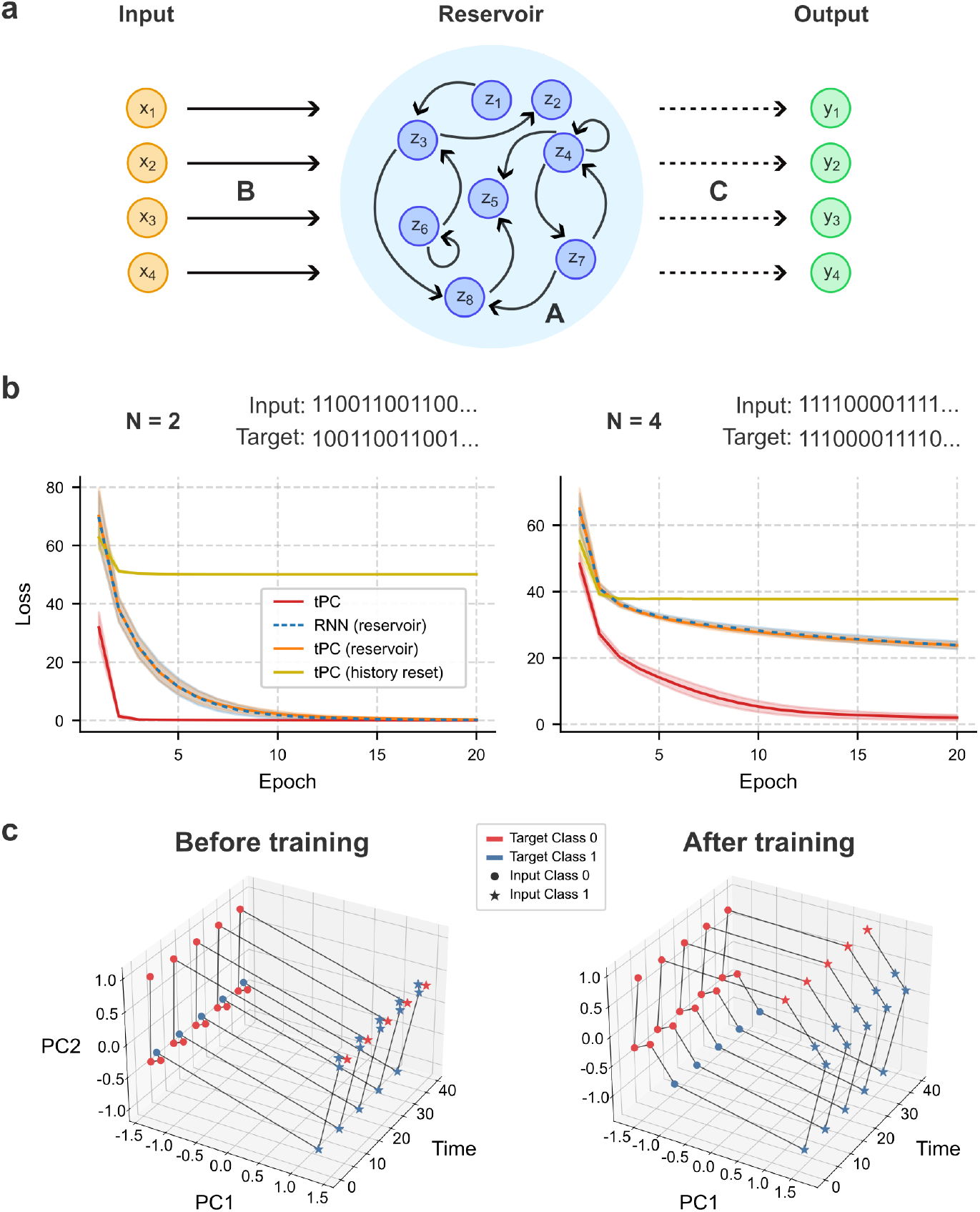
tPC shapes reservoir dynamics via local plasticity to facilitate downstream readout from the reservoir. **a:** The reservoir computing framework typically uses an RNN with fixed input and recurrent weights. Only the output weights are trained, as indicated by the dashed lines. **b:** Training loss curves for the alternating two-state task given a block length of either 2 or 4. tPC substantially improves performance over standard reservoir computing as the task becomes more difficult. **c:** Visualisation of tPC hidden state trajectories in the space of the first two principal components (PC1 and PC2) before and after training.

To demonstrate this, we consider an alternating two-state task, which consists of a sequence of alternating blocks of 0’s and 1’s (e.g., 110011001100…). The model performs next-step prediction: at each time step, it receives the current observation as input and is tasked with predicting the next observation. Given a block length of one, the sequence directly alternates between 0’s and 1’s and is therefore Markovian. As the block length is increased, the sequence becomes increasingly non-Markovian, such that longer temporal context is required to successfully predict the next observation. We include four models in our comparison: (1) standard reservoir computing implemented using an RNN with fixed input and recurrent weights; (2) an equivalent tPC formulation where again only the output weights were learned; (3) the standard tPC model, in which recurrent and input weights were also learned; and (4) a control tPC model in which the hidden state was reset at every time step, preventing information from propagating across time.

In the simplest non-Markovian setting, where the block length is set to two, both reservoir computing and tPC could successfully solve the task (Fig 3b). The RNN-based and tPC-based formulations of reservoir computing—distinguished only by the additional inference step in tPC—showed striking correspondence in performance. As expected, resetting the hidden state rendered the model unable to solve the task.

Importantly, as task difficulty increased, tPC increasingly outperformed reservoir computing (S2 Fig a). While the representations remain fixed with reservoir computing, tPC shapes neural trajectories in state-space during training to facilitate downstream decoding. This is also the case for instance when training RNNs using the influential FORCE algorithm [32] or its variants [33, 34], although tPC relies only on local plasticity.

To illustrate why plasticity at recurrent weights may be helpful, we projected the tPC hidden state onto its first two principal components in order to visualise its trajectory in time (Fig 3c). Prior to training, hidden states cluster according to the two possible input classes, with separation occurring primarily along PC1. Following training, however, the representations reorganise such that the two target classes become linearly separable based on PC1 and PC2. Together, these results demonstrate that tPC can exploit its reservoir dynamics to learn temporal dependencies over multiple time steps, while simultaneously shaping its internal representations through local plasticity in the recurrent weights.

### Learning complex cross-temporal interactions via a hierarchical architecture

Although the standard tPC model can learn simple non-Markovian sequences, we reasoned that such a framework may become limiting when learning more complex temporal dependencies. In contrast to tBPTT, which explicitly propagates error signals backward through time (Fig 4a), tPC relies on local updates and therefore learning depends more critically on the instantaneous hidden state representation to encode temporally relevant context. In task settings involving non-trivial cross-temporal dependencies, temporal structure may not be easily learnable from the current state representation.

**Fig 4.**
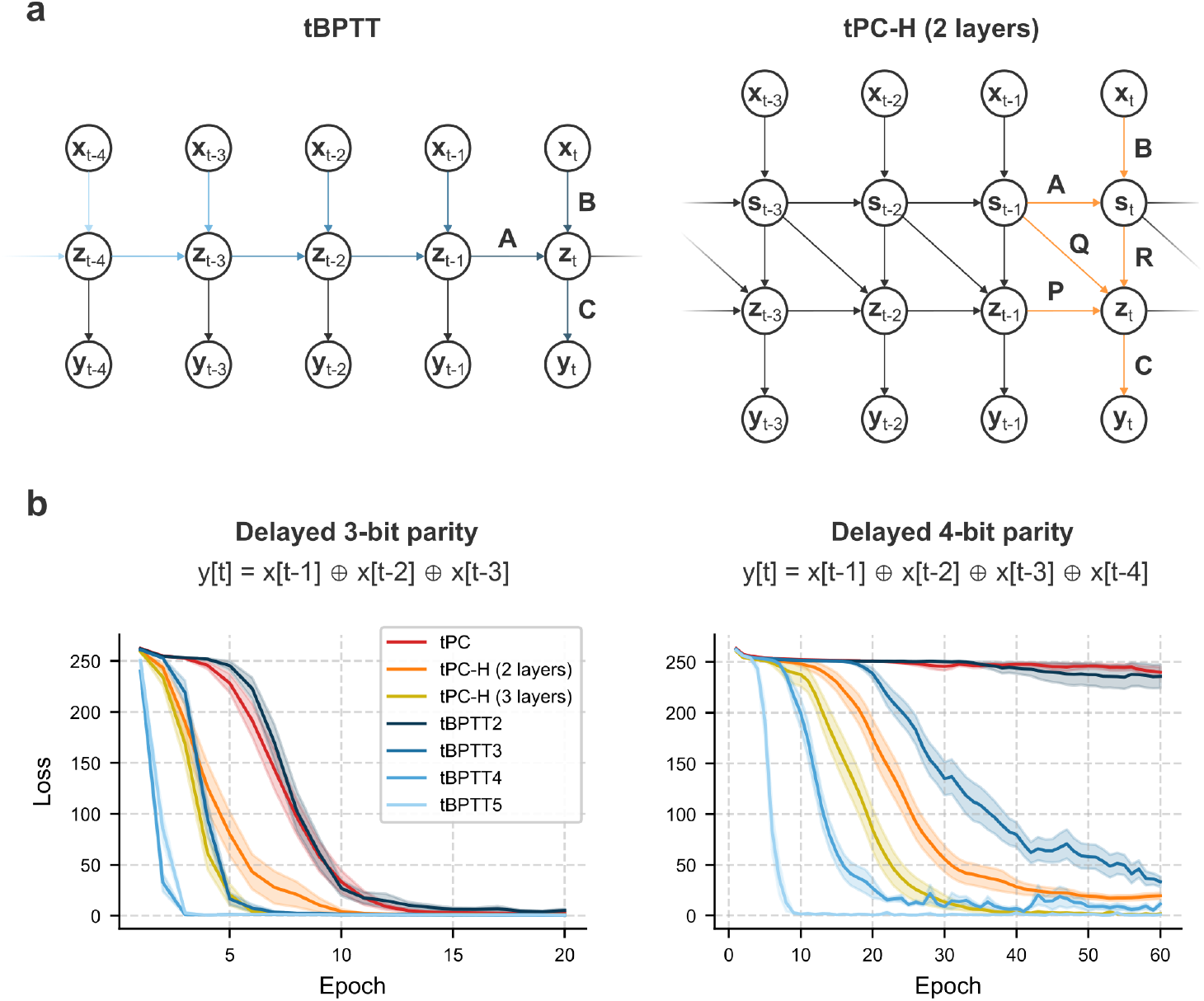
tPC-H supports learning of complex temporal dependencies through hierarchical recurrent dynamics. **a:** Comparison between tBPTT and tPC-H. On the left, the coloured arrows indicate which temporal influences are included when computing gradients of the loss using tBPTT, with progressively lighter shades of blue corresponding to longer truncation horizons. In tPC-H, the orange arrows instead indicate the local influences at each hierarchical levels that contribute to the gradient computation. **b:** Training loss curves for the delayed 3-bit and 4-bit parity tasks, highlighting how hierarchical recurrent dynamics may provide an alternative to tBPTT for learning more complex temporal dependencies.

We therefore hypothesised that more efficient learning of complex temporal dependencies could emerge from interactions between multiple recurrent reservoirs operating over partially distinct temporal representations. Intuitively, hierarchical recurrent dynamics may allow information to be integrated across multiple timescales while retaining local learning rules [35]. To make this idea more concrete, we implement a hierarchical version of tPC (tPC-H), where each layer receives as input both the current and previous hidden state from the preceding layer, thereby introducing an explicit temporal offset between hierarchical representations (Fig 4a; also see Fig 8). Each layer therefore maintains its own latent-state prediction error, resulting in an additional free-energy term for every hierarchical level. In the two-layer case, the free energy is given by:

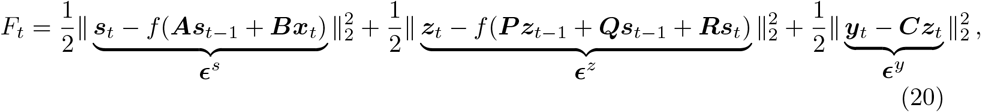

where inference should now be performed for each layer:

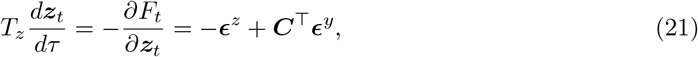

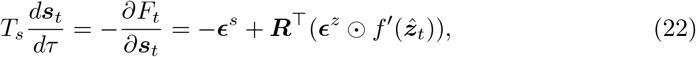

where we assume similar neuronal time constants *T*_*z*_ and *T*_*s*_ across layers for simplicity. Crucially, the weight updates, which may easily be extended to all synaptic connections, remain based on local Hebbian plasticity:

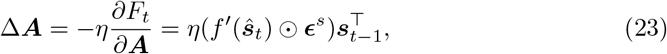

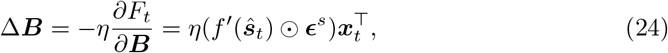

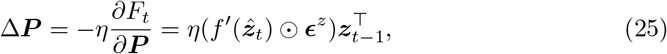

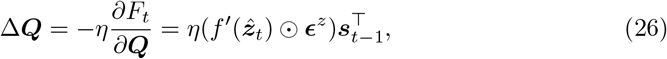

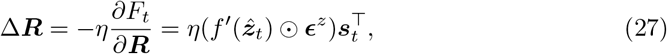

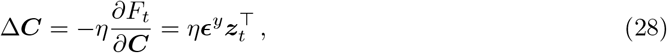

where ***ŝ***_*t*_ = ***As***_*t*−1_ + ***Bx***_*t*_ and 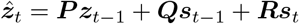. It is worth noting that we implement our model in discrete time here to facilitate comparison to standard RNN formulations. A more principled treatment of delays would likely require a continuous-time formulation, which lies beyond the scope of the present work.

To assess the performance of tPC-H, we turned to a delayed parity task, which requires the model to compute non-linear combinations of information distributed across distinct time steps. The task may be viewed as a temporal analogue of standard parity problems. Rather than processing simultaneous inputs, the network must maintain historical information over time and selectively integrate it to determine the final parity, producing an output of 1 if the cumulative number of 1’s in the input sequence is odd and 0 if it is even. Delayed parity tasks therefore provide a standard benchmark for evaluating a model’s ability to manage complex temporal dependencies.

On the delayed 3-bit parity task, all models—including tPC, tPC-H and tBPTT successfully learned the task (Fig 4b). As expected, increasing the truncation horizon for tBPTT improved learning efficiency. Notably, increasing the number of hierarchical layers similarly facilitated learning in tPC-H, with the 3-layer model closely matching the performance of tBPTT3. When task difficulty was increased, the single-layer tPC model was unable to solve the delayed 4-bit parity task. This is consistent with the hypothesis that decoding complex cross-temporal dependencies from a single reservoir-like representation becomes increasingly difficult as temporal structure grows more demanding. In contrast, both deeper tPC-H architectures and RNNs trained with longer tBPTT truncation horizons were able to solve the task successfully (S2 Fig b).

In addition to settings involving complex temporal dependencies, we reasoned that in the single-layer tPC model, memories of previous stimuli retained in the hidden state would also be vulnerable to subsequent distractors, particularly when these were of large amplitude. Since temporal context must remain recoverable from the evolving reservoir state, large perturbations to the hidden dynamics may interfere with downstream decoding. To test this, we consider a simple task in which the model was presented with one of two possible cues, followed by a distractor presented through the same input channel, before being required to reproduce the original cue upon a subsequent ‘Go’ signal (Fig 5a). Distractors were sampled from a Gaussian distribution with mean *µ* = 0 and varying standard deviation *σ*, which controlled distractor strength.

**Fig 5.**
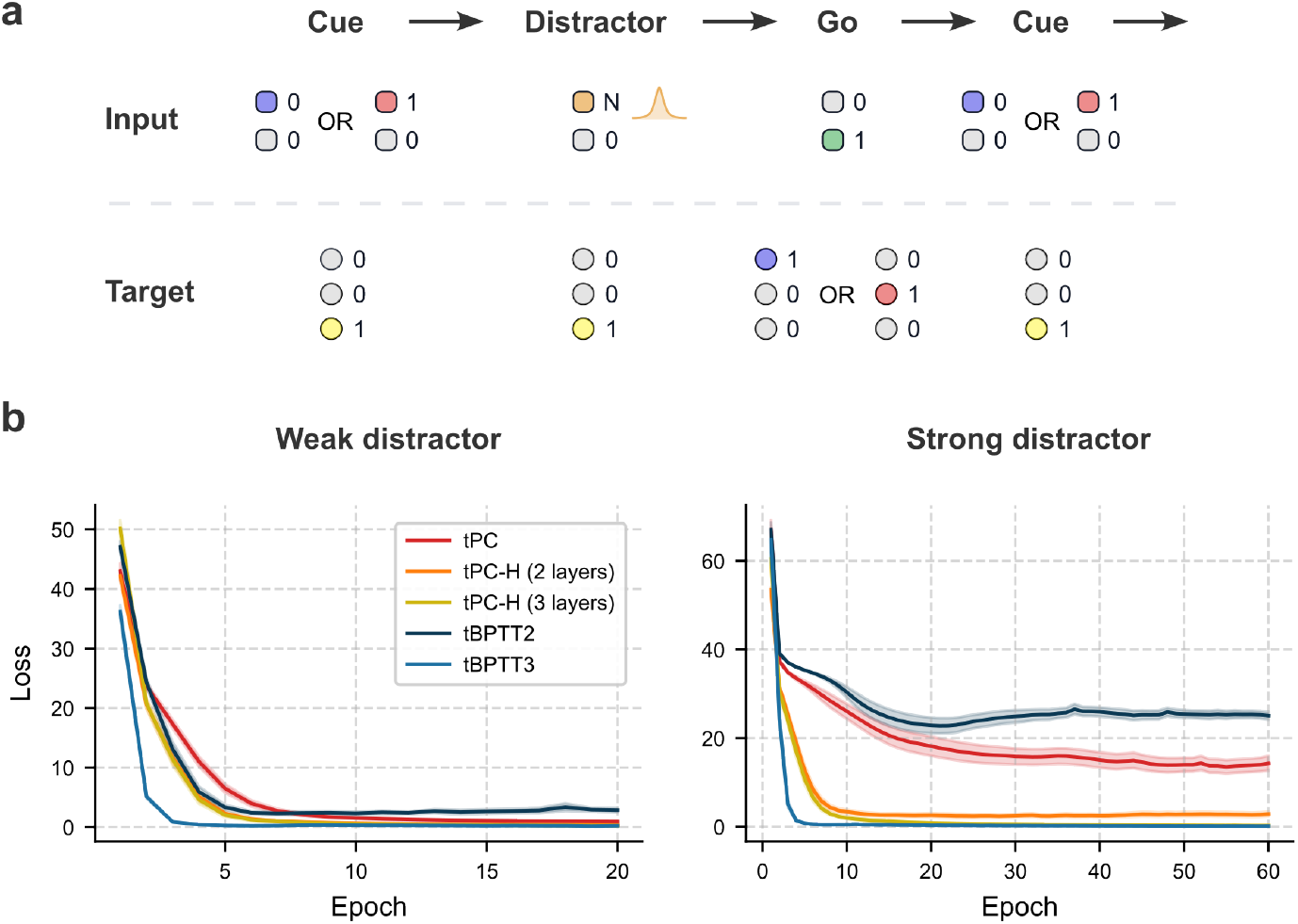
tPC-H confers robustness to strong distractors through selective temporal integration. **a:** Schematic of the distractor task, where the model is cued with one of two possible signals and then exposed to an intervening distractor before receiving a ‘Go’ prompt. **b:** Training loss curves for the distractor task, assuming either weak (*σ* = 1) or strong (*σ* = 10) distractors. As distractor strength was increased, the standard tPC model failed to reproduce the original cue.

When distractor strength was weak or comparable in magnitude to the cue input (*σ* = 1), the single-layer tPC model successfully reproduced the cue, with tPC-H providing only marginal improvement. However, under strong distractor conditions (*σ* = 10), tPC-H substantially outperformed single-layer tPC. This suggests that hierarchical recurrent pathways may enable the network to preserve task-relevant information by selectively integrating partially distinct temporal representations. For RNNs trained with tBPTT, successful performance required gradients to propagate back to the initial cue presentation. Importantly, we observed that the tPC-H architecture was robust to considerably larger perturbations (S2 Fig c). Together, these results demonstrate that tPC-H not only facilitates learning of complex temporal dependencies, but also confers robustness of information stored in the hidden state to distractors.

### Learning temporally extended context-dependent tasks via eligibility traces

Having shown that hierarchical recurrent dynamics enable efficient learning of complex temporal dependencies, we next investigate how tPC may be augmented with eligibility traces to support credit assignment across long temporal delays. Eligibility traces have been extensively studied in the context of three-factor learning rules and synaptic tagging and capture [36, 37]. Experimental evidence suggests that coincident pre- and post-synaptic activity does not necessarily induce immediate long-term potentiation (LTP). Instead, local co-activity can produce a transient biochemical state—commonly referred to as a synaptic tag or eligibility trace—which temporarily marks a synapse as eligible for subsequent plasticity [38]. Lasting synaptic modification may then occur only upon the later arrival of a global modulatory signal, such as dopamine [37].

Recent computational work has demonstrated how biologically inspired eligibility traces can be formalised in RNNs to support online learning without requiring BPTT. In particular, the Random Feedback Local Online (RFLO) algorithm [39] and the eligibility propagation (e-prop) framework [40] showed that RNNs can maintain local eligibility traces that accumulate historical activity and thereby enable temporally delayed credit assignment. Here, we extend this formulation to a predictive coding framework, yielding an augmented architecture which we refer to as tPC-E. Conceptually, tPC-E introduces a key modification to standard tPC: leaky recurrent dynamics, which allow inputs to directly influence activity over multiple time steps. To preserve accurate credit assignment, synaptic plasticity must be modified to match these activity dynamics.

Under local gradient approximation, this entails that synapses maintain exponentially decaying eligibility traces for weight updates. For clarity, we describe the linear case here and provide the non-linear formulation in the Methods (subsection “Derivation of tPC-E update rules”). For comprehensive treatments of the theory underlying such eligibility traces, we refer the reader to the original publications [39, 40].

Starting from the free energy:

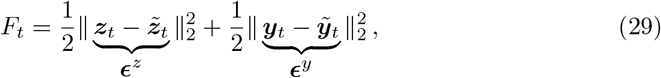

we modify the hidden-state dynamics to include a leaky integration term, where the timescale of temporal integration is governed by a parameter *α*:

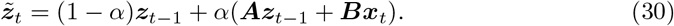

In standard tPC, updates to the recurrent and input weights depend only on the instantaneous activities ***z***_*t*−1_ and ***x***_*t*_, respectively (Eqs 5 and 6 restated for convenience, assuming no non-linearity):

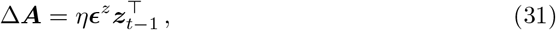

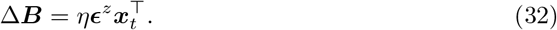

In tPC-E, however, the instantaneous activity terms 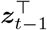 and 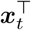 are replaced by exponentially decaying eligibility traces 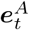 and 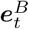:

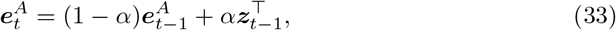

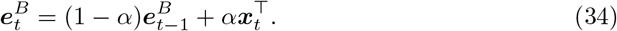

We show in the Methods (subsection “Derivation of tPC-E update rules”) that the following updates approximately minimise *F*_*t*_:

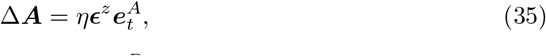

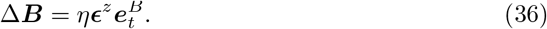

Intuitively, the eligibility traces allow synapses to retain a decaying memory of past activity, enabling temporally delayed error signals to influence learning across substantially longer timescales (Fig 6a).

**Fig 6.**
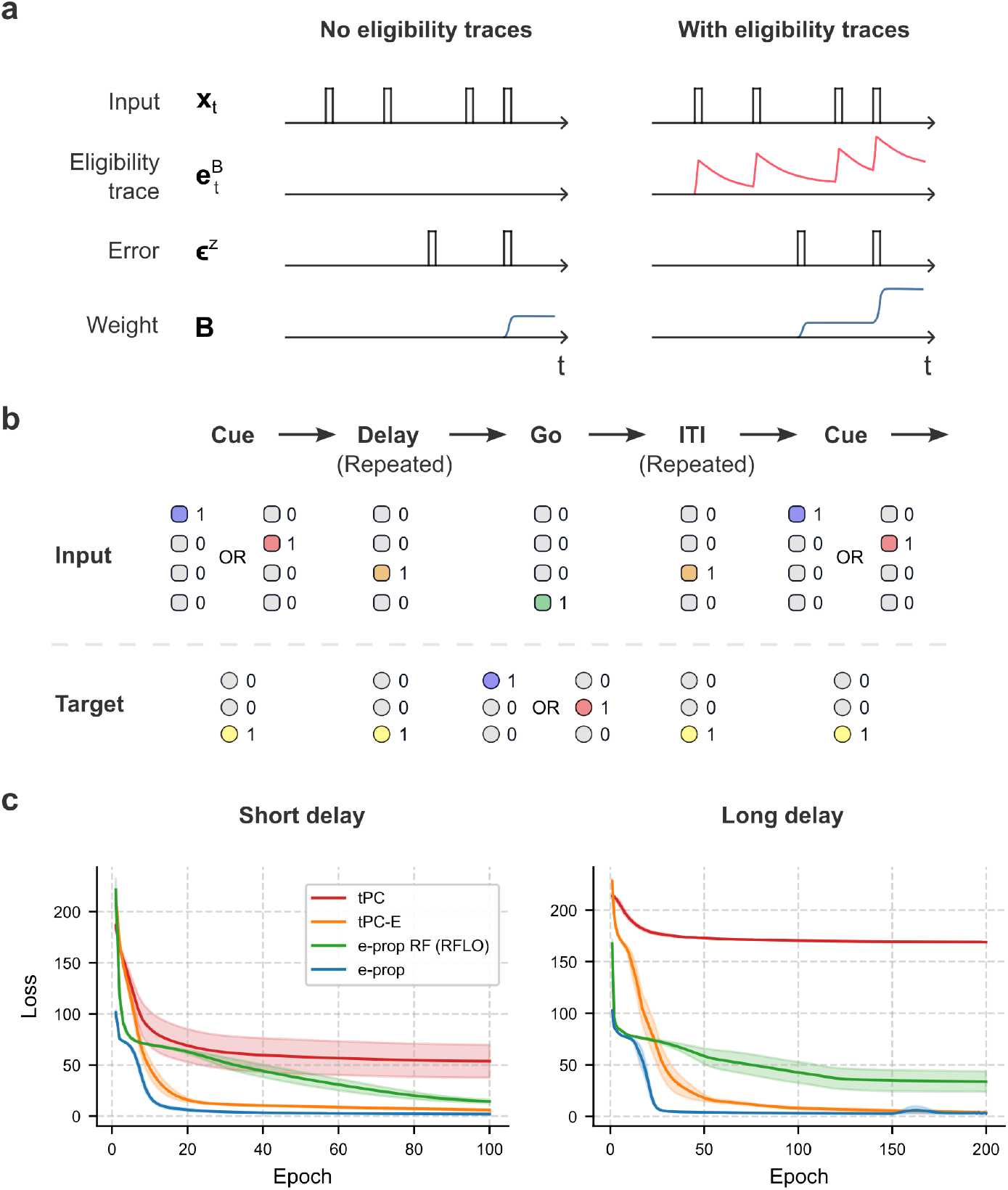
tPC-E enables long temporal gaps to be bridged via eligibility traces. **a:** In the absence of eligibility traces, Hebbian plasticity depends on the simultaneous activity of neurons encoding the input and the error signal. By contrast, eligibility traces preserve a transient synaptic memory of past activity, allowing later learning signals to drive plasticity. Hence in the presented example, the first prediction error in the right display triggers plasticity as the eligibility trace maintains memory of recent activity in the input. **b:** Schematic of the delayed response task, where each trial begins with the presentation of one of two possible cues, which the model had to reproduce after a delay of variable duration. Trials were separated by an inter-trial interval (ITI) of variable length (1 to 3 time steps). **c:** Training loss curves for the delayed response task, assuming either a short delay (1 to 5 time steps) or long delay (5 to 10 time steps). Eligibility traces were necessary for the model to recover the original cue even after short delays, although RFLO performance saturated as the delay length increased.

To demonstrate how eligibility traces may enable tPC to learn long-range temporal dependencies, we consider a working memory paradigm based on a delayed response task. At the beginning of each trial, the model was presented with an initial cue (0 or 1) and was required to reproduce the cue following a delay period of variable length (Fig 6b). Trials were separated by an inter-trial interval (ITI) of variable duration. In addition to standard tPC and the proposed tPC-E model, we compared against RNN-based implementations of both e-prop and RFLO. Although e-prop was originally introduced in the context of spiking neural networks, we implemented it here using a rate-based RNN with forward state dynamics given by

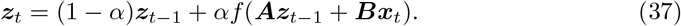

RFLO refers here to an analogous formulation in which fixed random feedback weights are introduced to maintain biologically plausible local learning rules. While the original publication introduced several versions of e-prop, we consider here its basic implementation, which may be viewed as equivalent to RFLO with the feedback weights set equal to the readout weights (see Methods, subsection “e-prop and RFLO implementation”).

Without eligibility traces, standard tPC failed to reliably reproduce the cue under both short and long delay conditions (Fig 6c). In contrast, both tPC-E and e-prop successfully solved the task even under extended delay conditions, although e-prop converged more rapidly during training. RFLO was able to handle relatively short delays, but performance did not extend to longer delay durations (S2 Fig d). These findings demonstrate how eligibility traces may support temporally delayed credit assignment within predictive coding networks. More broadly, they show that tPC-E can facilitate learning across extended temporal gaps while retaining update rules that remain local in both space and time.

## Discussion

BPTT is generally acknowledged to be biologically implausible, but remains widely used in computational neuroscience. This is due to its effectiveness in learning complex temporal dependencies that are central to many behavioural and neural processes, including sequence generation, sensory prediction, working memory, and decision-making. Here, we presented a framework for extending predictive coding networks to learn such temporal dependencies while relying only on local synaptic plasticity.

After first establishing a structural equivalence between the weight updates for tPC and tBPTT1, we showed that tPC can exploit reservoir dynamics to encode short-range temporal context, while simultaneously shaping neural trajectories in state space to support downstream readout. We next demonstrated that introducing a hierarchical architecture with delays facilitates learning of non-trivial temporal dependencies and enables robust maintenance of memory content even in the presence of strong distractors. Finally, we showed that tPC networks can be augmented with biologically inspired eligibility traces to solve tasks involving long delay durations. Together, these findings position tPC within a broader landscape of temporal learning frameworks (Fig. 7), highlighting important connections to reservoir computing, BPTT, and e-prop, in addition to the Kalman filter correspondence identified in previous work [21].

**Fig 7.**
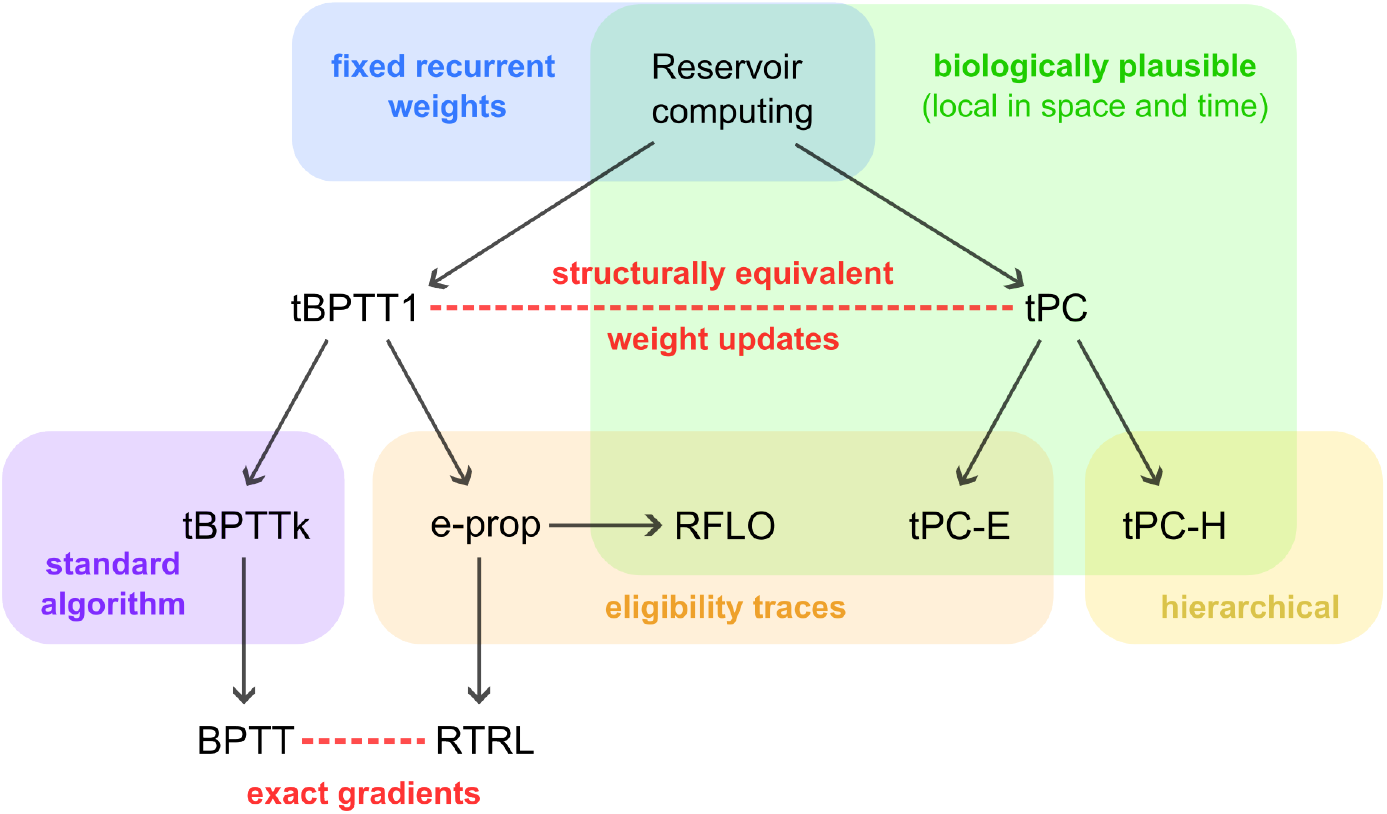
Relationship between different algorithms. tPC is closely related to several established frameworks for learning temporal dependencies. In this work, we showed that tPC is structurally similar to tBPTT1, both of which extend the reservoir computing framework by introducing plasticity at the input and recurrent weights. tPC can also be combined with eligibility traces or hierarchical recurrent dynamics, giving rise to tPC-E and tPC-H, respectively. Eligibility traces had previously been explored in RNNs through eligibility propagation (e-prop), which can be made more biologically plausible by incorporating random feedback weights (RFLO). More generally, tBPTT remains the standard training procedure for RNNs, using a truncated backward pass of length k. In contrast to the aforementioned algorithms, BPTT and RTRL compute exact gradients by propagating error through the full sequence or by accounting for all possible dependencies, respectively. Arrows denote modifications from one model to the next, with top-to-bottom progression indicating an improvement in prediction accuracy over the preceding model.

We aimed here to identify and investigate various computational ingredients that biological neural networks may exploit to support temporal learning across different settings, which we explored here using relatively simple tasks. Our work therefore invites several natural extensions. First, while our discrete-time implementation of tPC-H captures delayed interactions at a functional level, it does not provide a fully satisfactory treatment of delays from the perspective of predictive processing theory [35], which may instead call for a continuous-time formulation. In the case of tPC-E, we assumed a fixed inverse time constant of *α* = 0.5, consistent with previous work [41]. A more biologically principled approach would, however, consider a distribution of time constants, consistent with the substantial heterogeneity observed across neural systems [42, 43]. Eligibility traces could also be incorporated across multiple hierarchical levels, potentially with distinct temporal dynamics at each level. Finally, future work should test whether these mechanisms scale to richer behavioural settings involving higher-dimensional inputs and more complex temporal dependencies. Recent work on how dendritic morphology and neuronal dynamics may support temporal learning in deep networks provides one possible direction for such extensions [44].

While we focused our comparison here on BPTT, another influential algorithm, real-time recurrent learning (RTRL) [45], also computes exact gradients of the loss function with respect to recurrent weights by accounting for dependencies across all time steps. Unlike BPTT, RTRL performs these computations online, but at the cost of prohibitive computational and memory requirements, in addition to relying on non-local update rules. Nevertheless, the appeal of online gradient computation has motivated numerous efforts to derive tractable approximations to RTRL [41]. RFLO represents one such constrained approximation, enforcing locality within the weight update rule to obtain biologically plausible learning dynamics [39]. More recently, promising progress has been made towards closer approximations of RTRL by appealing to neuromodulatory mechanisms [46, 47]. An alternative approach, which has also recently been extended to tPC [48], albeit with unclear biological grounding, is to apply RTRL to recurrent architectures with element-wise recurrence and complex-valued weight matrices, yielding substantial reductions in computational complexity [49].

A central focus of this work has been on how biologically plausible mechanisms may support learning of temporal structure over relatively short timescales. At longer timescales, however, temporal credit assignment remains challenging even for RNNs trained with BPTT due to the vanishing and exploding gradient problem, whereby error signals propagated over many recurrent steps either decay or grow unstably [27, 28]. Gated recurrent architectures, such as long short-term memory networks (LSTMs) and gated recurrent units (GRUs), were introduced in part to address these limitations by preserving information over extended temporal intervals through learned gating mechanisms [50, 51]. Other approaches—including Neural Turing Machines [52], memory networks [53], and related memory-augmented architectures [54, 55]—have equipped neural networks with explicit memory systems that support the storage and retrieval of information beyond the hidden state of a conventional RNN. Together, these developments suggest that long-timescale learning may depend not only on improved temporal credit assignment algorithms, but also on architectural mechanisms that support memory maintenance and flexible retrieval. The mechanisms studied here should therefore be viewed as addressing one part of the temporal learning problem, rather than as a complete account of long-timescale memory and credit assignment.

Of note, although biological plausibility is framed here primarily in terms of locality in space and time, a deeper understanding of neural computation could substantially broaden the scope of biologically plausible temporal credit assignment. In the case of BPTT, for example, hippocampal reverse replay [56, 57] raises the intriguing possibility that biological systems may possess mechanisms for propagating information backward across experienced temporal sequences [10]. Moreover, recent studies have found evidence that different stages of a behavioural sequence may be represented by distinct but systematically related neural populations in prefrontal cortex [58, 59]—mapping temporal sequences onto a spatial pattern of activity. Such representations could, in principle, provide a substrate through which error-related signals are assigned across time, potentially enabling richer forms of temporal credit assignment than those captured by standard locality-based formulations. Finally, we also acknowledge that the biological plausibility of the Adam optimiser may be debated. Importantly, however, its use does not compromise the locality of the tPC learning rule, since weight updates are independently rescaled based on synapse-specific state variables. We further note that individual synapses may plausibly maintain multiple internal states [60], and recent work has suggested potential biological implementations of Adam-like optimisation [61].

Despite recent advances in machine learning, state-of-the-art architectures such as transformers often rely on increasingly long context windows to capture long-range dependencies, incurring substantial memory and computational costs. Understanding how biological systems achieve efficient temporal credit assignment may therefore offer useful insights for both neuroscience and machine learning. Progress towards such an understanding will likely depend on close interaction between experimental and theoretical neuroscience, both to uncover novel biological mechanisms and to formalise their computational principles.

## Methods

### Derivation of BPTT update rules

Although the update rule for BPTT is well-established in the literature, we include the derivation for the recurrent weight matrix ***A*** (Eq 13) here for completeness and clarity in the context of our model. Assuming similar forward dynamics (Eq 12), we consider the loss for a sequence of length *T* :

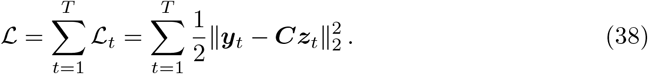

We derive the gradient descent update direction for ***A***, setting the learning rate to 1 here for simplicity,

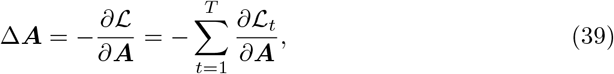

For a single time step, the chain rule gives

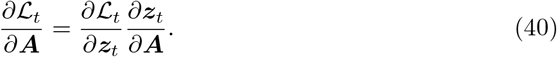

The first term is simply

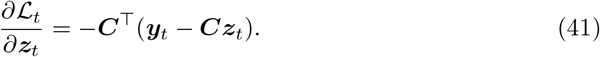

The second term, 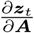, requires a more careful treatment. Since the same recurrent weight matrix ***A*** is used at every time step, ***z***_*t*_ depends on ***A*** both directly, through the transition at time *t*, and indirectly, through the dependence of previous hidden states on ***A***. Thus,

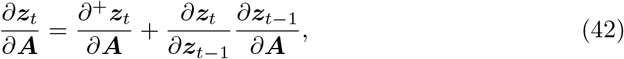

where 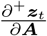 denotes the immediate partial derivative of ***z***_*t*_ with respect to ***A***, treating ***z***_*t*−1_ as fixed. Recursively expanding the second term gives

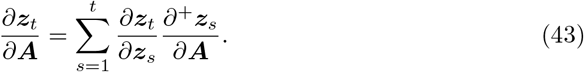

Since 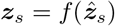 and 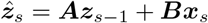, we can write the immediate derivative as

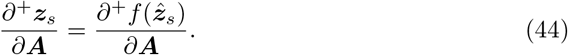

Substituting into the expression for the gradient and taking the negative-gradient direction yields

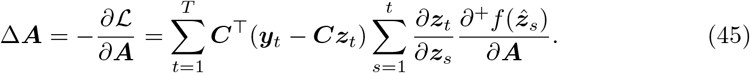

This is the full BPTT update direction for the recurrent weights.

The Jacobian relating the hidden state at time *t* to the hidden state at an earlier time *s* can be expressed as a product of local recurrent Jacobians along the trajectory:

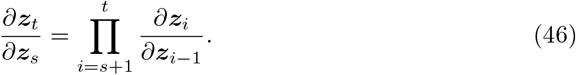

For the recurrent dynamics considered above, each local Jacobian is given by

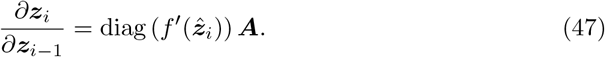

Consequently, the gradients propagated by BPTT involve repeated multiplication by recurrent Jacobians.

### Derivations of tPC update rules and fixed point

Here we derive the tPC update rule for the recurrent weights ***A***, corresponding to Eq 16, and the fixed point of the tPC inference dynamics in the linear case, corresponding to Eq 18.

For convenience, we first rewrite the local tPC objective at time *t* (Eq 3):

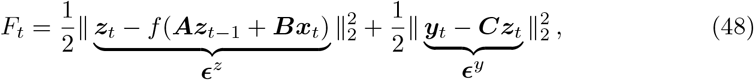

The corresponding inference dynamics update the hidden state ***z***_*t*_ in the direction that reduces *F*_*t*_:

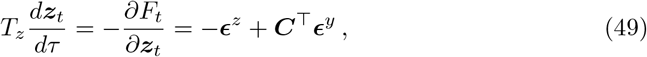

At convergence of the inference dynamics, 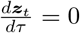, and therefore

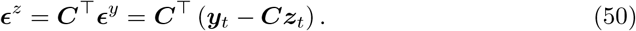

Following convergence, the recurrent weights are updated by gradient descent on *F*_*t*_. Assuming a learning rate of 1 again for simplicity, and treating the inferred activity ***z***_*t*_ as fixed during the weight update, we have

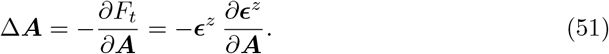

Since 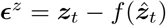 and 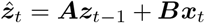, the second term is given by

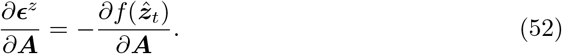

Substituting this expression and the fixed-point condition from Eq 50 into the weight update equation gives

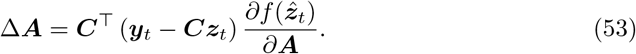

Since tPC performs this update locally at each time step, the total update over a sequence is obtained by summing over time:

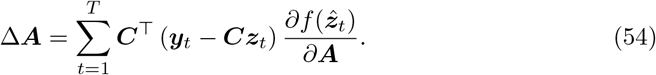

This is the tPC recurrent weight update given in Eq 16. It has the same form as the one-step truncated BPTT update, except that the hidden states ***z***_*t*_ are obtained after tPC inference rather than by a purely feedforward recurrent pass. We note that Eq 52 implicitly treats the previously inferred hidden state ***z***_*t*−1_ as fixed with respect to ***A***, and therefore does not yield the exact gradient of the tPC energy with respect to the weights. In principle, the exact gradient could instead be obtained by differentiating through the dependence of previous hidden states on ***A*** using BPTT. However, this would introduce non-local dependencies across time, compromising the temporal locality and biological plausibility of the tPC learning rule.

We next derive the fixed point of the tPC inference dynamics in the linear case. Starting from the fixed-point condition in Eq 50, but this time taking *f* to be the identity function,

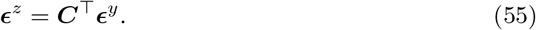

Expanding both prediction errors gives

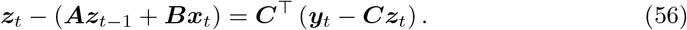

Rearranging terms yields

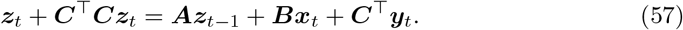

Therefore,

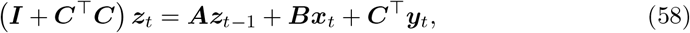

and hence, we end up with

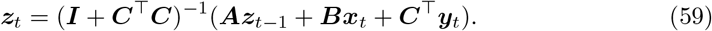

This is the linear fixed point of the tPC inference dynamics given in Eq 18.

### Update rules for 3-layer tPC-H model

Similar to the two-layer case, each layer of receives as input both the current and previous hidden state from the preceding layer (Fig 8), such that the free energy is given by:

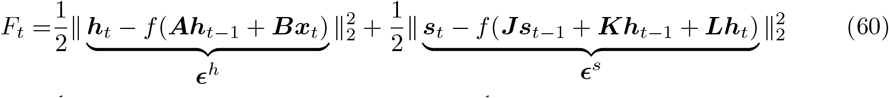

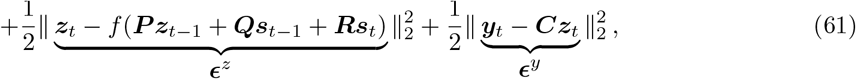

where inference is performed for each hierarchical layer:

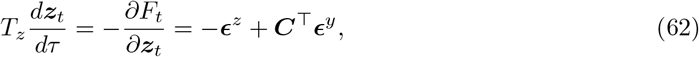

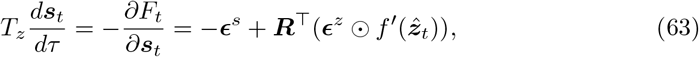

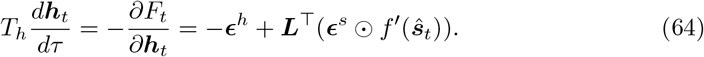

The full list of weight update equations corresponding to all synaptic connections is thus:

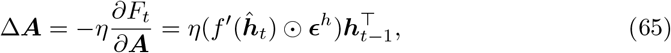

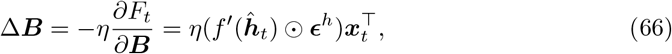

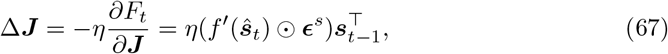

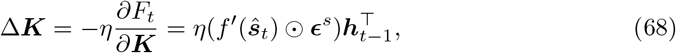

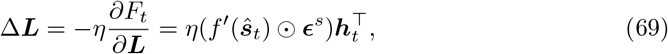

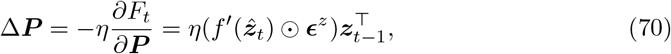

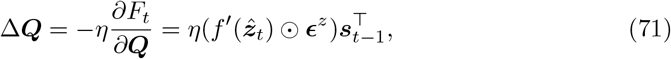

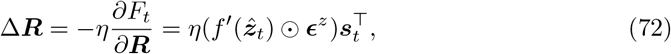

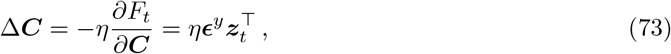

Where 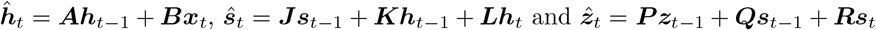.

**Fig 8.**
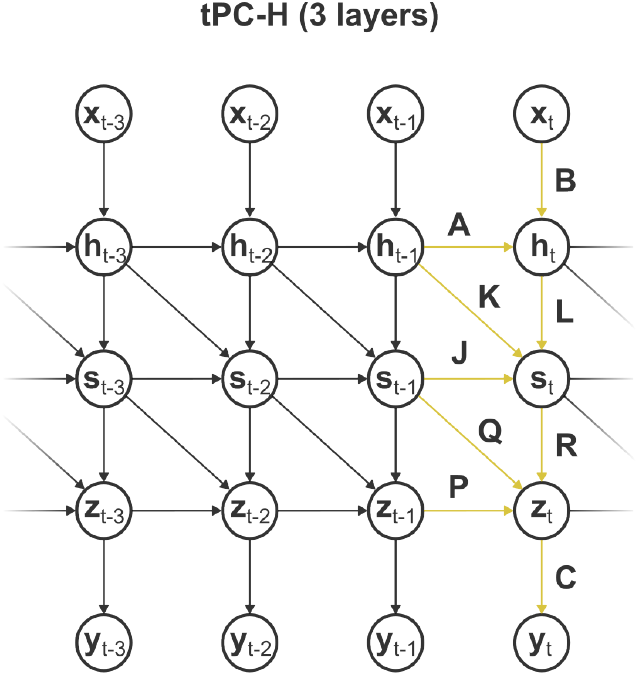
3-layer tPC-H model. 3-layer variant of tPC-H architecture presented in Fig 4a. ***x***_*t*_ and ***y***_*t*_ denote the input and observation, respectively, while ***h***_*t*_, ***s***_*t*_ and ***z***_*t*_ represent the three hierarchical hidden layers. The yellow arrows highlight the local influences at all three hierarchical levels that contribute to the gradient computation.

### Derivation of tPC-E update rules

The mathematical foundation of online eligibility traces in RNNs has been rigorously established through two distinct methodological lenses [39, 40]. Murray [39] truncated the exact Real-Time Recurrent Learning (RTRL) framework for RNNs—omitting spatially non-local terms and incorporating random feedback weights—to arrive at the RFLO algorithm. Bellec et al. [40] developed the e-prop algorithm by rewriting the loss gradient in a factored form that reveals the maximum amount of gradient information accessible through local online computation. Here, drawing from the principle of free-energy minimisation, we describe how the eligibility trace formulation proposed by e-prop and RFLO may be extended to the tPC framework.

If we return to the free energy

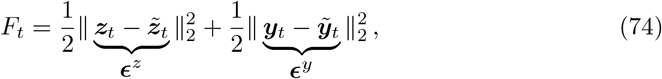

and include a leaky integration term in the hidden state dynamics

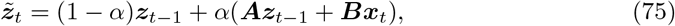

while allowing for a non-linear readout of the hidden state

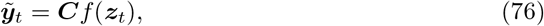

we can expand out the free energy to obtain

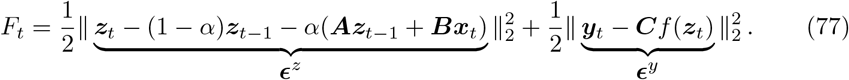

The inference dynamics are now given by

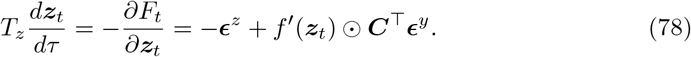

As in standard tPC, to find the updates of the recurrent and input weights, we take the negative of the derivative of the free energy with respect to the weights as follows:

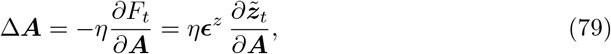

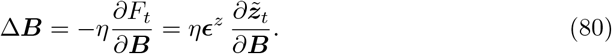

Suppressing neuron- and synapse-specific indices for notational simplicity, the gradient of the predicted dynamics 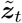 over recurrent weights ***A*** is equal to:

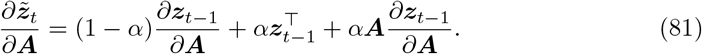

Here, similar to e-prop and RFLO, we ignore the last term in the above equation as its computation would not be local. Additionally, we approximate the gradient of ***z***_*t*−1_ with the gradient of 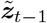, enabling us to replace the instantaneous gradient term in the left hand side of Eq 81 with a leaky eligibility trace that accumulates recent synaptic activity over time:

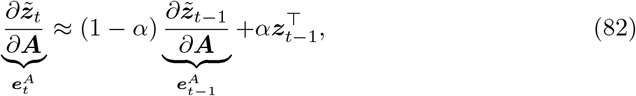

such that the updates for the recurrent weights are given by

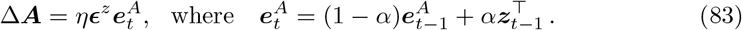

Analogous approximation for the input weights gives the following update:

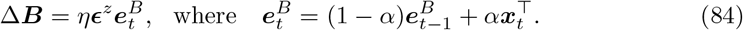

We can see that when eligibility traces are absent (i.e., *α* = 1), the weight updates reduce to those of standard tPC (assuming no non-linearity):

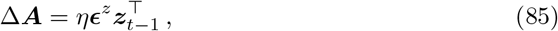

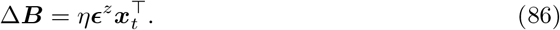

The updates for the output weights are similar to those in the standard tPC:

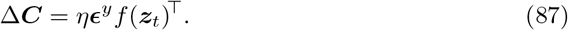

It is worth noting that, in our implementation of tPC-E, we found it advantageous to place the non-linearity in the observation model rather than in the hidden state dynamics. Acknowledging that this modification results in hidden state dynamics that do not exactly match those of the non-linear tPC formulation used in earlier sections of the paper when *α* = 1, our primary objective here was to introduce eligibility traces into the tPC framework, in direct analogy to e-prop and RFLO. We therefore adopt this formulation as a practical means of studying the effect of eligibility traces.

### e-prop and RFLO implementation

Here, we briefly describe the rate-based implementations of e-prop [40] and RFLO [39] used in this paper. To keep the presentation concise, we use compact matrix notation and again omit the neuron- and synapse-specific indices commonly used in related derivations.

For both algorithms, the network dynamics are given by

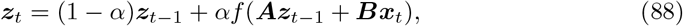

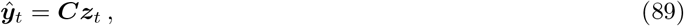

with output prediction error

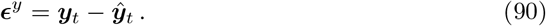

Following the local approximation used here, the recurrent and input eligibility traces are updated according to

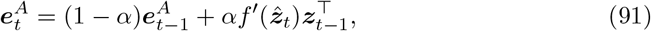

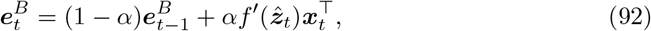

where

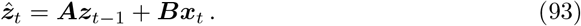

For e-prop, the output prediction error is projected back to the recurrent units using the transpose of the readout weights, giving

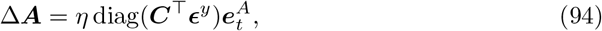

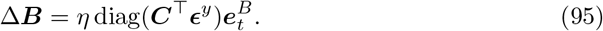

Here, diag(·) forms a diagonal matrix from the neuron-specific learning signals, which scale the corresponding rows of the eligibility matrices.

For RFLO, ***C***^⊤^ is replaced by a fixed random feedback matrix ***F***, whose entries are independently sampled from 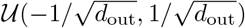, where *d*_out_ is the output dimension:

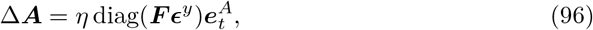

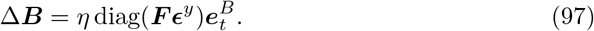

Thus, in the implementations considered here, e-prop and RFLO use the same eligibility traces and differ only in whether symmetric or random feedback is used to construct the learning signal.

The output weights are updated directly from the output prediction error for both algorithms:

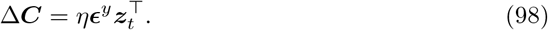

### Model training

All models were implemented in PyTorch [62] and were trained online with a batch size of 1. Network weights were initialised using Kaiming uniform initialisation [63], and all layers included bias terms. To facilitate comparison, parameter counts were matched across the models shown in Fig. 4 and Fig. 5. Hyperparameters were selected by grid search over learning rate and inference step size. Each result is reported as the mean across at least five random seeds, with standard errors computed across seeds. The full set of hyperparameters used for each task is reported in Tables 1, 2 and 3. For tPC, the loss was recorded at each time step from the observations predicted by Eqs 9 and 10, prior to inference and therefore before the target observation was made available to the model. This ensured that predictive performance was evaluated under the same constraints as for tBPTT-trained RNNs.

**Table 1.** Hyperparameters for Fig 2 and Fig 3.

| **Hyperparameter** | **Fig 2** | **Fig 3** |
|---|---|---|
| Number of hidden units | 32 | 32 |
| Learning rate (*η*) | 0.00005 | 0.0005 |
| Inference step size (1/*T*<sub>*z*</sub>) | 0.01 | 0.1 |
| Number of inference steps | 20 | 20 |
| Batch size | 1 | 1 |
| Epochs | 100 | 20 |
| Number of seeds | 5 | 5 |

**Table 2.**
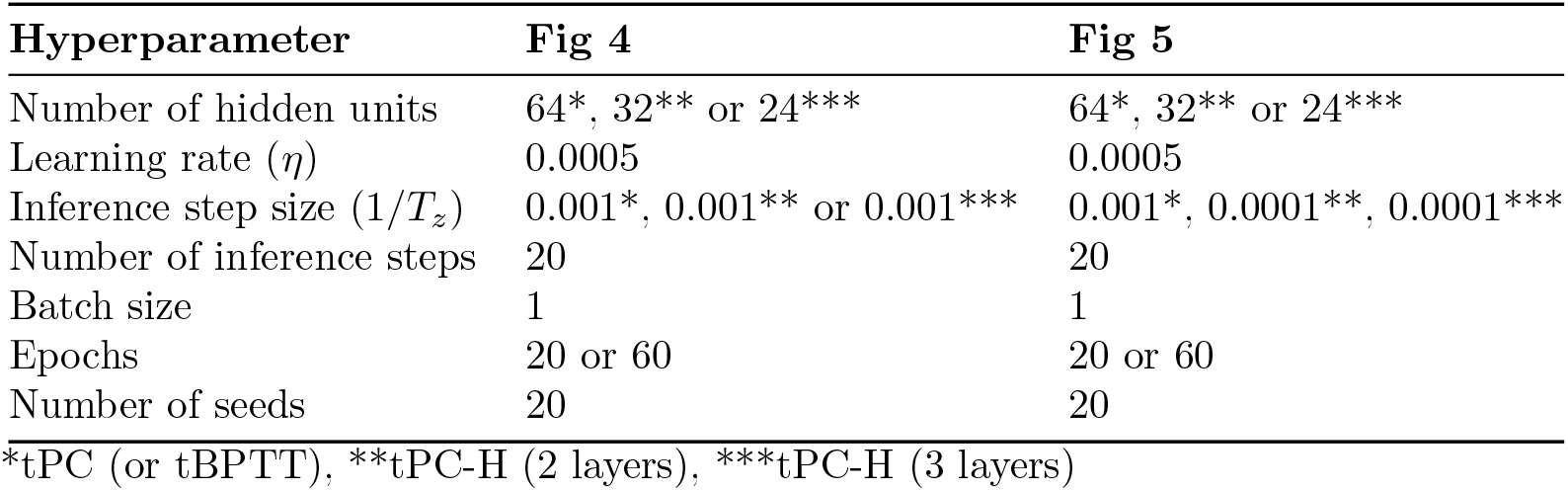
Hyperparameters for Fig 4 and Fig 5.

| **Hyperparameter** | **Fig 4** | **Fig 5** |
|---|---|---|
| Number of hidden units | 64<sup>*</sup>, 32<sup>**</sup> or 24<sup>***</sup> | 64<sup>*</sup>, 32<sup>**</sup> or 24<sup>***</sup> |
| Learning rate (*η*) | 0.0005 | 0.0005 |
| Inference step size (1/*T<sub>z</sub>*) | 0.001<sup>*</sup>, 0.001<sup>**</sup> or 0.001<sup>***</sup> | 0.001<sup>*</sup>, 0.0001<sup>**</sup>, 0.0001<sup>***</sup> |
| Number of inference steps | 20 | 20 |
| Batch size | 1 | 1 |
| Epochs | 20 or 60 | 20 or 60 |
| Number of seeds | 20 | 20 |

<sup>\*</sup>tPC (or tBPTT), <sup>\*\*</sup>tPC-H (2 layers), <sup>\*\*\*</sup>tPC-H (3 layers)

**Table 3.** Hyperparameters for Fig 6.

| **Hyperparameter** | **Short delay** | **Long delay** |
|---|---|---|
| Number of hidden units | 32 | 64 |
| Learning rate (η) | 0.05* | 0.05* |
| Inference step size (1/*T*<sub>*z*</sub>) | 0.01 | 0.01 |
| Number of inference steps | 20 | 20 |
| Leak parameter (α) | 0.5 | 0.5 |
| Batch size | 1 | 1 |
| Epochs | 100 | 200 |
| Number of seeds | 5 | 5 |

\*The learning rate for RFLO was reduced to 0.005 to improve training stability.

Models were trained using the Adam optimiser, except for the results in Fig 6, for which the eligibility-trace learning rule was applied directly. For experiments using Adam, the optimiser adaptively rescales parameter updates to improve numerical efficiency and stability [64]. Adam maintains exponential moving averages of the gradient 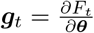 and its element-wise square 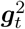,

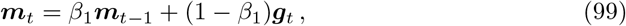

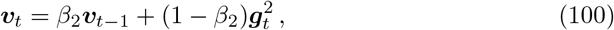

such that the parameters are updated according to

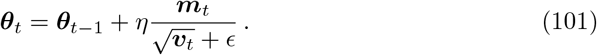

We used the default values *β*_1_ = 0.9 and *β*_2_ = 0.999, with *ϵ* = 10^−8^ included for numerical stability. We have omitted the bias correction step here for simplicity. Although Adam introduces a separate pair of scalar states for each parameter, it preserves the parameter-wise locality of the underlying tPC learning rule: the Adam update for a given parameter depends only on its corresponding local learning signal and its own first- and second-moment estimates, without requiring learning signals associated with other parameters.

## Acknowledgments

The authors would like to acknowledge the use of the University of Oxford Advanced Research Computing (ARC) facility in carrying out this work (https://doi.org/10.5281/zenodo.22558).

## Supporting information captions

**S1 Fig. Similar performance between tPC-P and RNN-P on the position estimation task**. Results corresponding to Fig 2, but now also including tPC-P. tPC-P, which receives position information both implicitly via the inference process, and explicitly from the previous observation, performed similarly to RNN-P on the position estimation task.

**S2 Fig. Task performance across different difficulty levels**. Minimum loss as a function of task difficulty for the tasks considered in **a:** Fig 3 (alternating two-state task), **b:** Fig 4 (delayed parity task), **c:** Fig 5 (distractor task), and **d:** Fig 6 (delayed response task). Hyperparameters were the same as those reported in Tables 1, 2 and 3 (Long delay), and were not separately optimised for each difficulty level. In **d**, the delay duration was fixed here rather than variable, and the ITI was set to a single time step. Results are based on 5, 10, 10, and 3 random seeds for **a**-**d**, respectively.

**S1 Fig.**
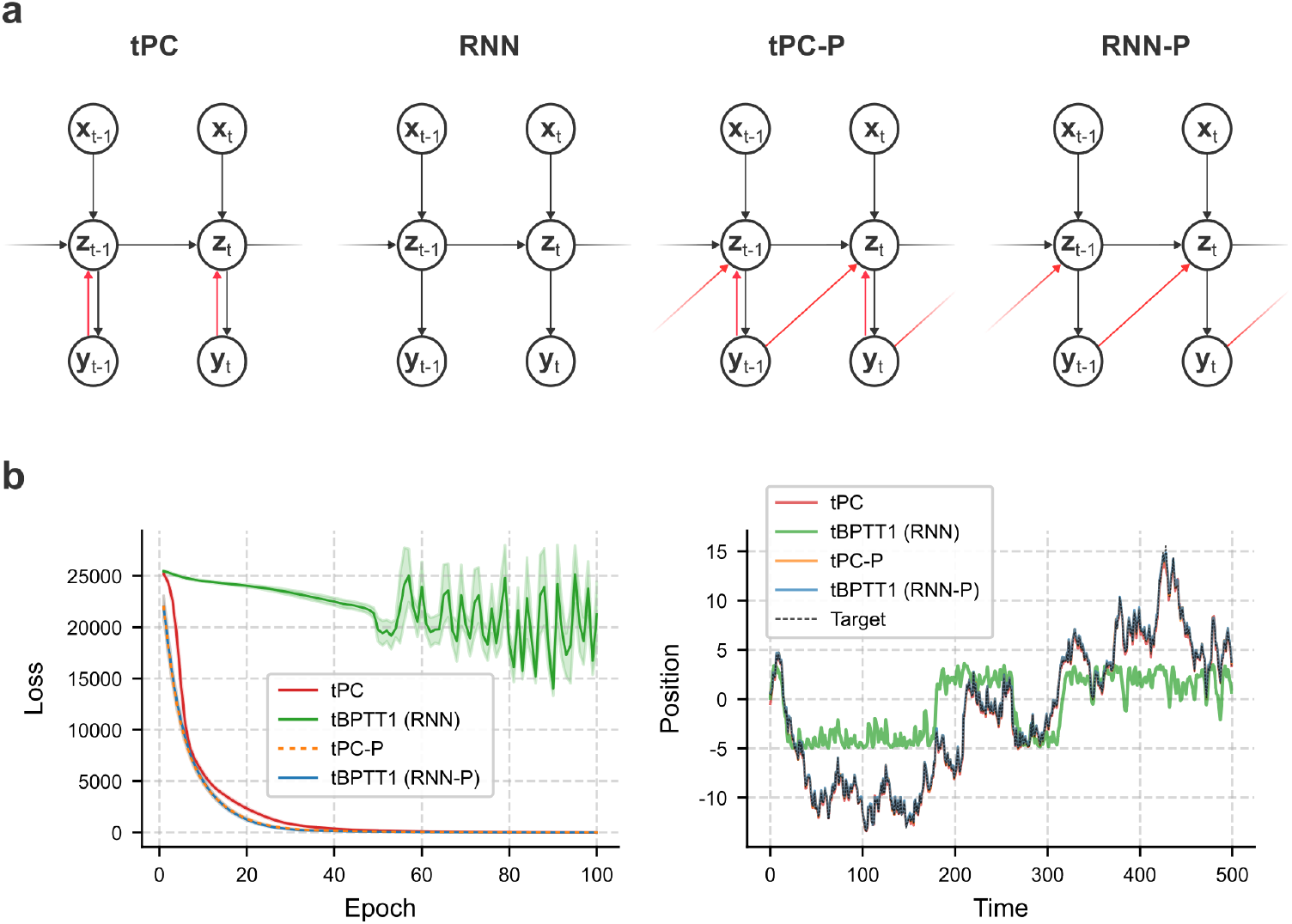
Similar performance between tPC-P and RNN-P on the position estimation task. Results corresponding to Fig 2, but now also including tPC-P. tPC-P, which receives position information both implicitly via the inference process, and explicitly from the previous observation, performed similarly to RNN-P on the position estimation task.

**S2 Fig.**
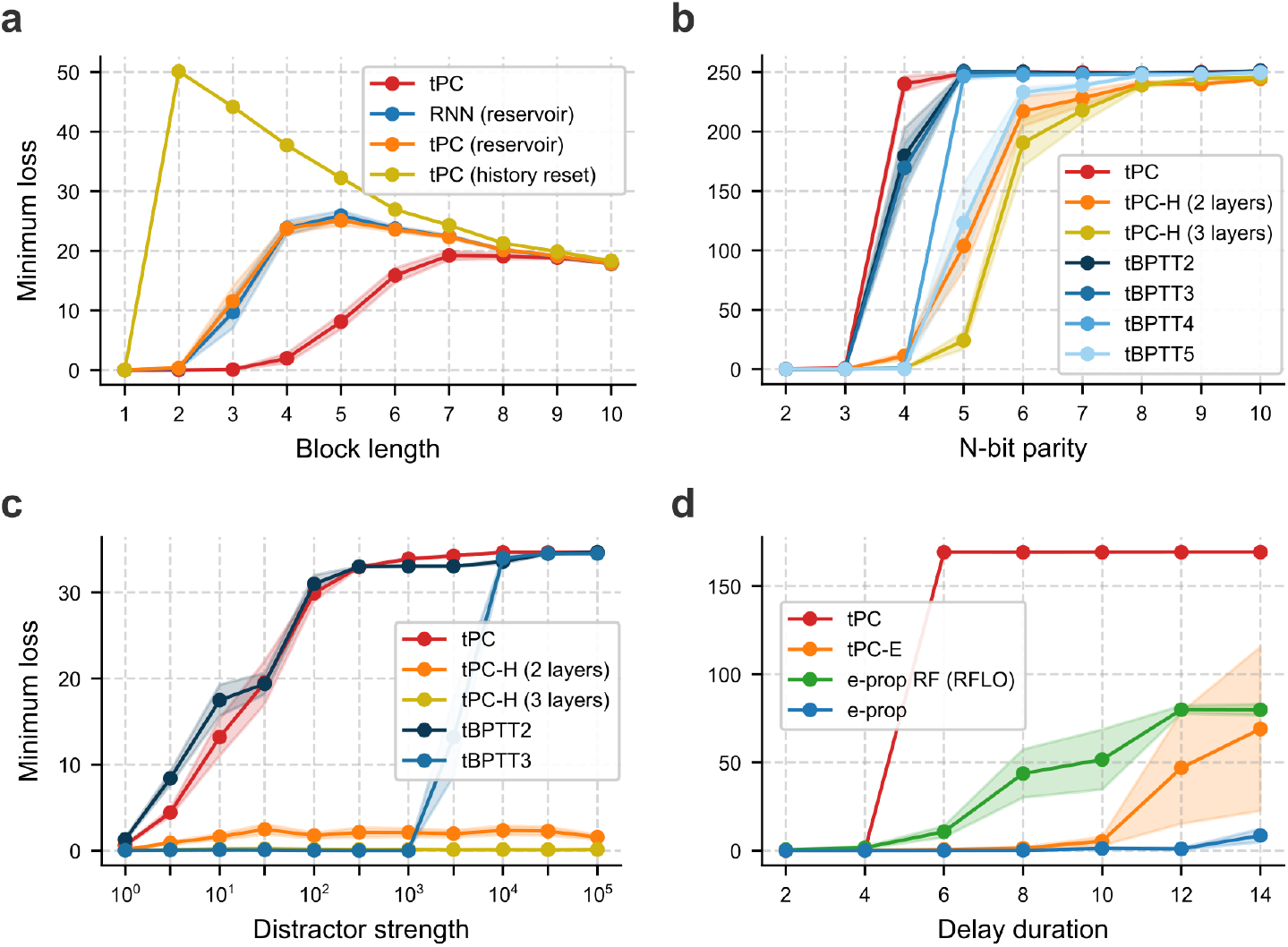
Task performance across different difficulty levels. Minimum loss as a function of task difficulty for the tasks considered in **a:** Fig 3 (alternating two-state task), **b:** Fig 4 (delayed parity task), **c:** Fig 5 (distractor task), and **d:** Fig 6 (delayed response task). Hyperparameters were the same as those reported in Tables 1, 2 and 3 (‘Long delay’), and were not separately optimised for each difficulty level. In **d**, the delay duration was fixed instead of variable, and the ITI was set to a single time step. Results are based on 5, 10, 10, and 3 random seeds for **a**–**d**, respectively.

